## Supplementary figures and images for "Kelpwatch: A new visualization and analysis tool to explore kelp canopy dynamics reveals variable response to and recovery from marine heatwaves"

### Supplemental Figure 1

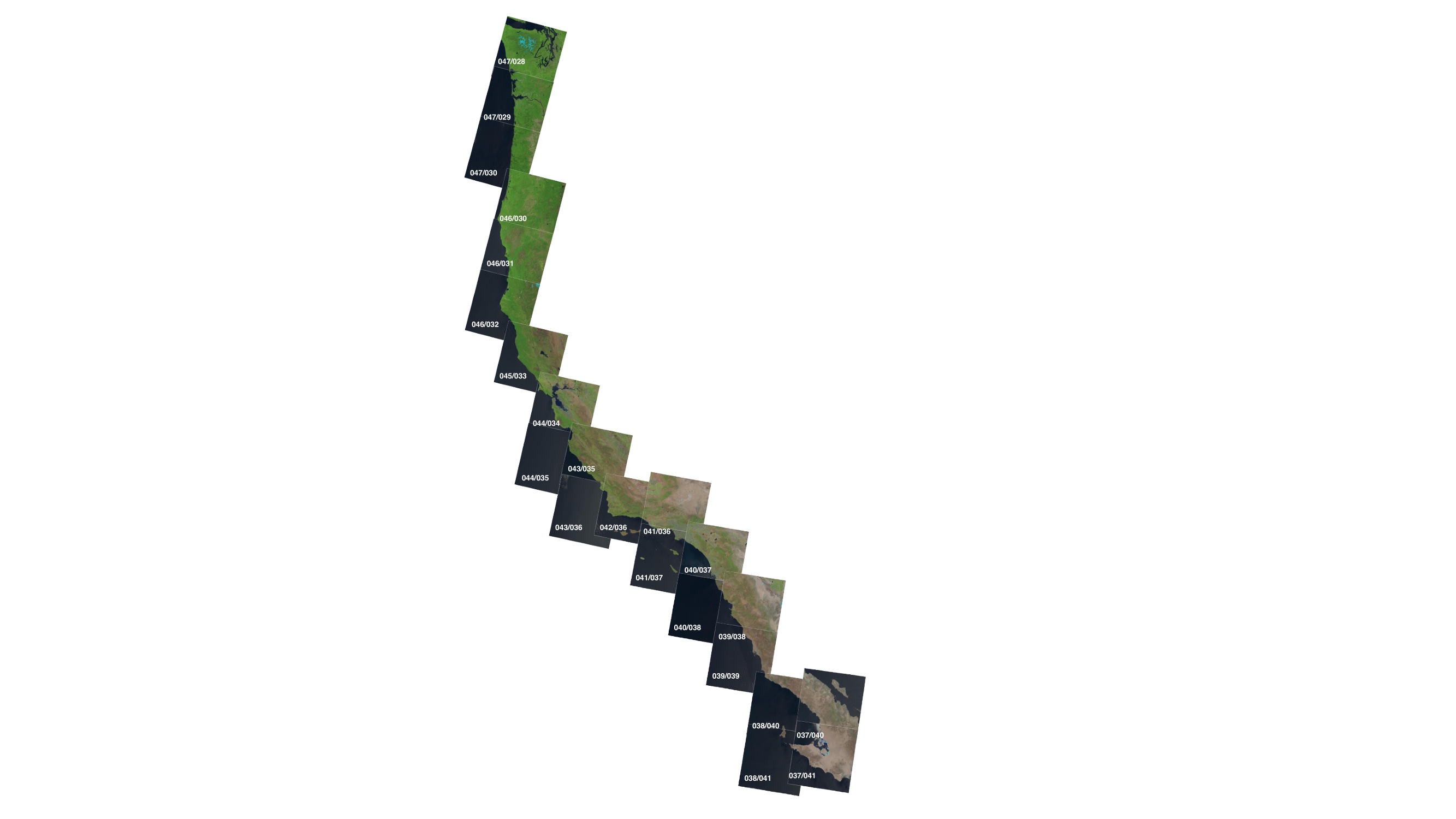

### Supplemental Figure 2

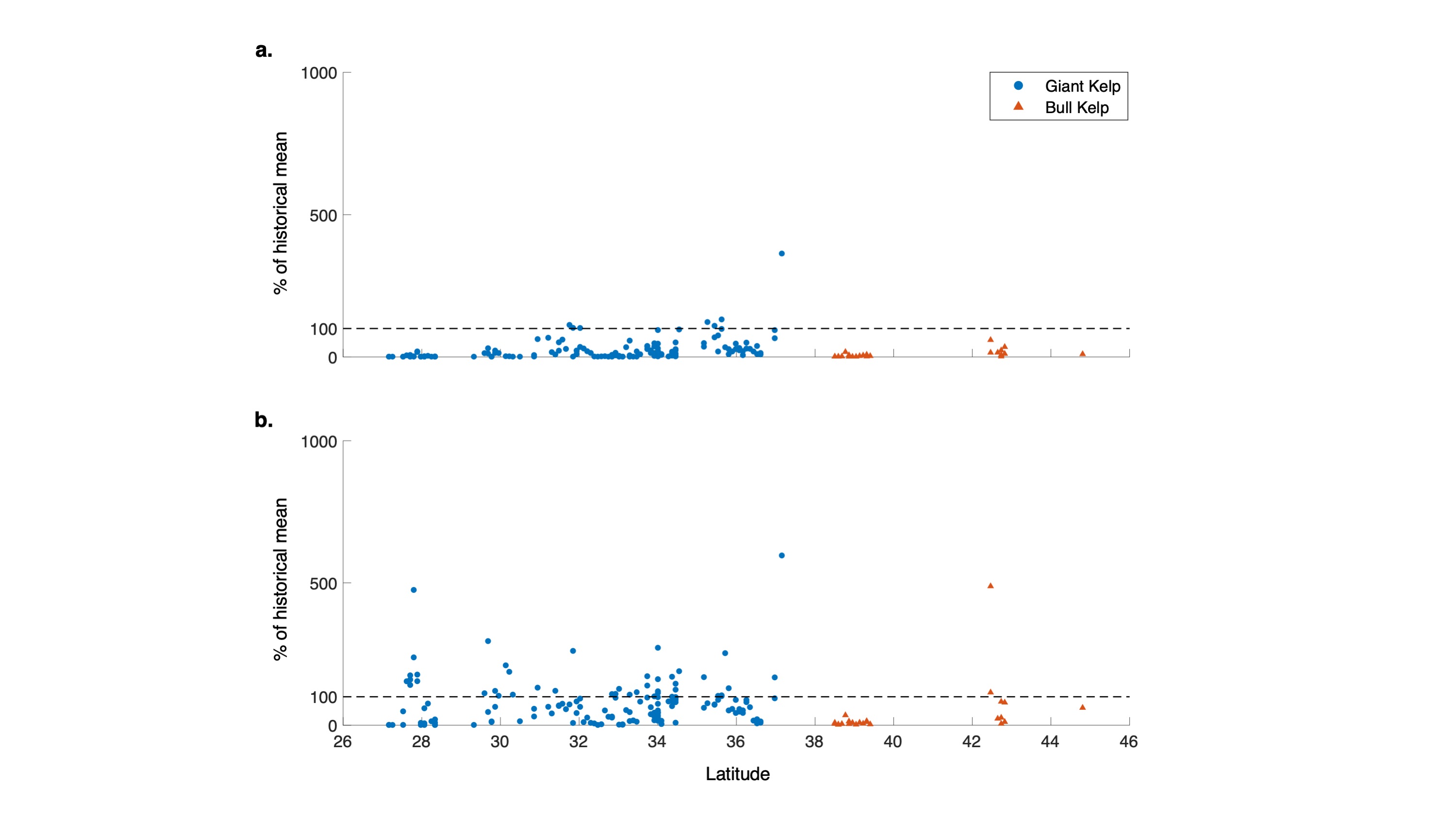

### Supplemental Table 1

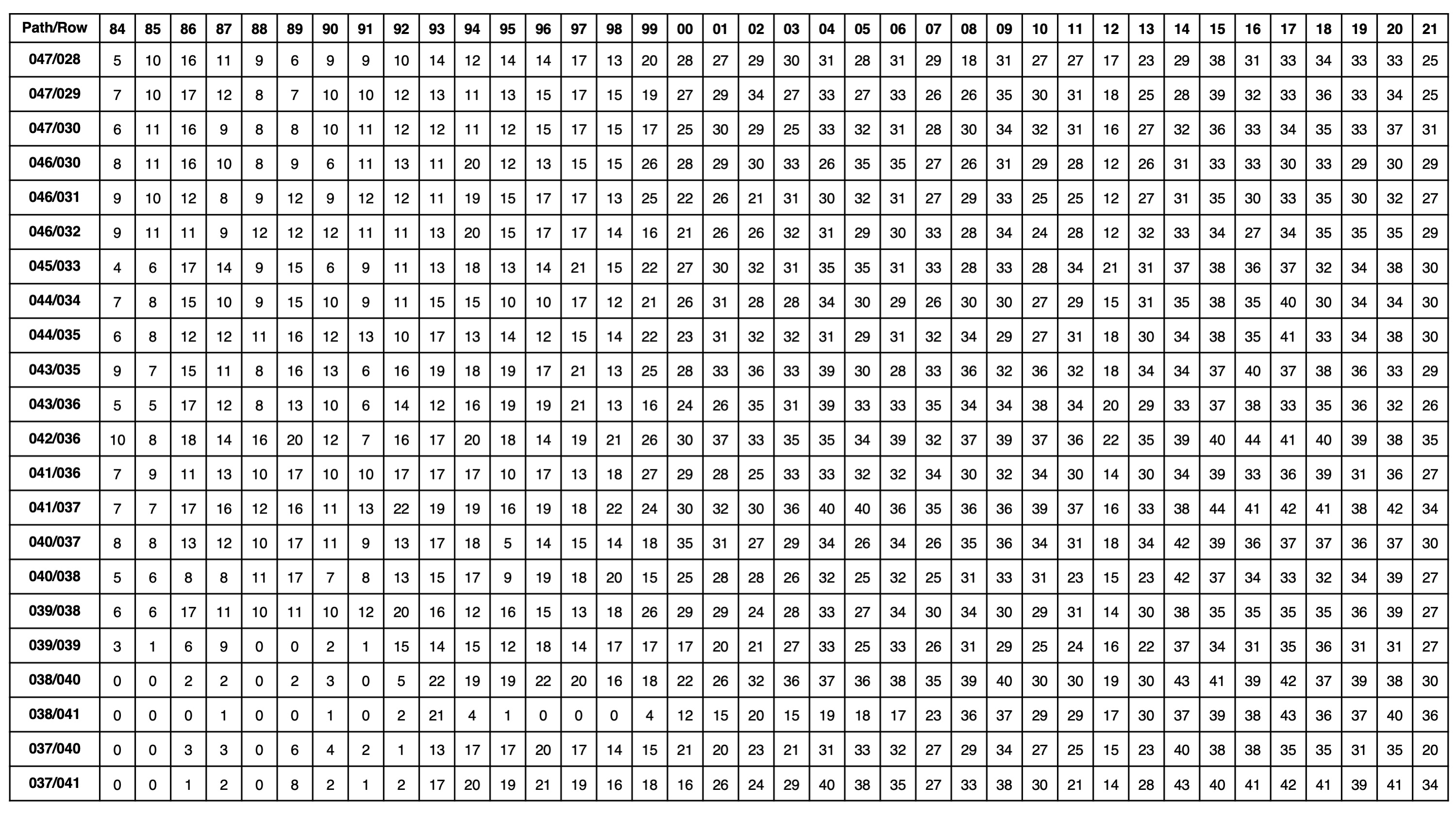
